# A transition to a more efficient attentional strategy facilitates motor learning in the presence and absence of movement-evoked experimental pain. A cross-sectional experimental study

**DOI:** 10.64898/2026.06.03.729955

**Authors:** David Matthews, Ali Khatibi, Deborah Falla

## Abstract

Pain demands attention and can disrupt task-related goals. Attention allocation is a key cognitive process supporting motor learning and disruption of internal schemas associated with attentional control during motor learning can result in interference in improvements in performance. Movement-contingent pain is an important characteristic of persistent musculoskeletal pain. Despite this, research exploring pain interference with motor learning and attention has exclusively utilised tonic pain paradigms. Understanding the impacts of movement-contingent pain on motor learning and attention may provide important insights into the interaction between pain and motor learning. The aim of this study was to; 1) explore the robustness of a movement-contingent pain paradigm across an extended period of training, 2) explore the impact of movement-contingent pain on improvements in performance and attentional allocation during motor learning.

Three groups (healthy non-pain, healthy experimental-pain and persistent pain experimental-pain) completed ten trials of a motor sequence learning task while experiencing a movement-contingent electrical stimulation. Three task performance measures and five gaze indices, previously associated with attentional control, were collected. Results showed that; 1) low frequency electro-cutaneous stimulation could produce a valid and consistent pain experience across a sustained period of training, 2) attentional allocation becomes more efficient across learning, accompanied by improvements in task performance, 3) changes in task performance and attentional measures across training were similar in all groups despite the presence of pain, 4) movement-contingent experimental pain enhanced spatial performance at all time points in healthy participants but was not accompanied by a different pattern of attentional allocation.

This study demonstrates that the impact of movement-contingent pain on motor learning is comparable to the impacts of tonic experimental pain and provides interesting insights into patterns of attentional allocation across time but little evidence that these attentional allocations are impacted by the presence of pain or a past history of pain.

## 1. Introduction

The ability to caress a ball with a foot to generate the perfect flight and speed to score a goal or, manipulate a musical instrument with hands to create the desired intonation and sense of rhythm to grab attention provides indisputable evidence the human body has an unparallelled propensity to adapt to perform complex goal-orientated movements. This ability to create ever more effective and successful movements is the essence of motor learning [1]. Despite the importance of learning in everyday life, Lechner, Squire and Byrne (1999, p.79) stated *‘the experience of everyday life shows that perseverative tendencies of different parts of a train of thought can be weakened considerably by turning one’s attention with energy to a different matter’*

Attentional resources are considered finite [2]. Therefore, how attentional resources are allocated during motor learning influences behavioural improvements [3] and the propensity for formation of new memories. Eye tracking technology has been used in research as an indirect measure of attention to understand attentional allocation during motor learning [4–6]. A transition from high attentional requirements to low attentional requirements [7] and from feedback orientated to task orientated attentional strategies [4–6] has been observed across the stages of motor learning.

Pain demands attention [8] and therefore has the potential to interfere with these attention patterns observed during motor learning but increasingly evidence is indicating that pain can also focus attention [9–13]. Inherent to the many early theories around pain and attention, is that a salient stimulus, such as pain, automatically grabs attention interfering with any other tasks being completed. As the focus on theories around pain has moved away from the constrained idea of pain being about ‘detection of damage’, to a focus on the subjective lived experience of an individual centred around perceived threat [14], theories around attention are following a similar path. Legrain and colleagues in 2009, proposed the neurocognitive model of attention which suggested attentional allocation was a result of a balance between bottom-up (unintentional) and top-down (intentional) processes [15]. The Attention Schema Model [16] builds on this, suggesting a mechanism whereby higher centres are involved in both the involuntary (internal attentional schema) and voluntarily (conscious reallocation) control of attention, enabling both proactive and reactive strategies.

Experimental pain models have been used extensively in translatory pain research [17] including research exploring the impact of pain on motor learning [18]. Tonic pain paradigms have been the predominant pain paradigms used to explore interference with motor learning and attention as they are associated with less motor adaptation than movement evoked phasic pain paradigms [19], and therefore provide a better method to explore this interaction. That said musculoskeletal pain, characterised by pain evoked by movement, is a common aspect of persistent pain [20]. Meulders (2020) reported that a feature of persistent pain is the transition from pain being associated with a movement, to pain being triggered by the movement. The resultant effect is that the painful movement becomes an integral part of the pain experience. As tonic pain is considered only one aspect of musculoskeletal pain it therefore may only provide part of the picture of sensorimotor and attentional adaptations to pain during learning [22]. In addition, there remain questions around how meaningful a tonic pain experience is, when it is not associated with everyday living. Tabor et al, (2020) suggest that pain interference should be viewed in the context of motivation. A lack of perceived value placed on a painful experience may negate the need to interfere with goal orientated tasks. Studies exploring the impact of clinical pain on motor learning [23–26] have attempted to overcome the lack of ecological relevance of experimental pain experiences, but a lack of data on, or reports of low pain levels during motor learning tasks have made these studies difficult to interpret. Gallina et al, (2021) proposed a method of using low frequency sinusoidal electrical stimulation to induce a reliable short-term movement-contingent pain to the knee, modulated by the amount of quadriceps contraction. This method of producing movement-contingent experimental pain has the potential to overcome the above challenges and provide interesting insights into pain interference with motor learning and attention.

The study aims to investigate 1) the robustness and face validity of the movement-contingent experimental pain paradigm in creating a consistent pain experience across multiple trials, 2) the impact of the movement-contingent experimental pain on performance across motor learning, 3) the validity of the gaze indices to characterise attention during the motor learning, and 4) the impact of the movement-contingent experimental pain on the pattern of attentional allocation.

## 2. Methods

### 2.1. Participants

Seventy-seven right-handed participants were recruited from the student population at the University of Birmingham, United Kingdom, between 01/06/2023 and 31/05/2025. A priori power analysis (G*power) calculated a sample size of 55, based on using a repeated measures design, 80% power, an alpha of 0.05 and an expected small effect size (Cohen 0.25). A small effect size of 0.2 [27] was chosen to maximise power of the study as there was a lack of previous data on both the motor learning task performance and gaze indices used in this study.

Sixty-two healthy participants were randomized into either a control group or intervention group. Fifteen participants with a history of persistent (>3 months) pain (S1 Table) were recruited to the pain-intervention group. Potential participants were excluded from the study if they had any history of neurological, psychiatric or acute musculoskeletal disease or had non-corrected issues with vision. All groups were exposed to either a non-noxious or noxious electrical stimulation during motor learning, depending on their group allocation.

The cross-sectional experimental study received written ethical approval from the University of Birmingham ethics committee (ERN_21-0738A). Data from eleven participants was excluded from all data analysis processes either due to not completing the experimental protocol or unexpected deviations from the methodology (S2 Table). A further sixteen participants were excluded from analysis of the eye gaze data due to the eye gaze data not meeting the required level of quality across all trials. The characteristics of the participants excluded did not significantly differ from the other groups.

### 2.2. Experimental Protocol

Participants attend a lab at the University of Birmingham for a single session of motor learning training. On arrival Participants read the participant information sheet and provided written informed consent prior to starting the study. Demographic data was collected and participants completed The Fear of Pain Questionnaire (FOP-III) [28], The Pain Catastrophizing Scale (PCS) [29], The Depression, Anxiety and Stress Scale (DASS) [30] and The Attention Control Scale (ACS) [31]. The experimental procedure is outlined in figure 1.

**Fig 1:**
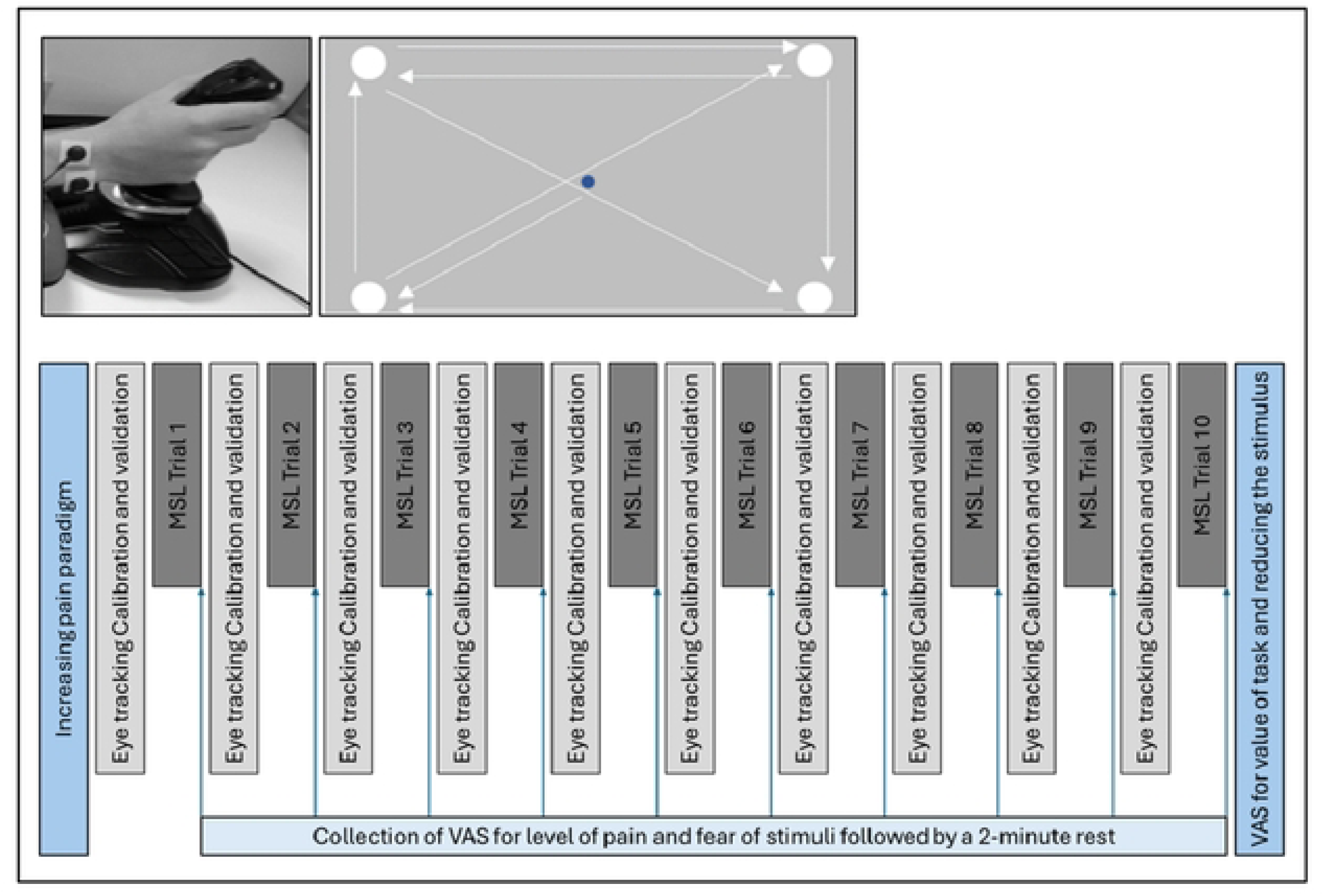
Experimental procedure. MSL – Motor sequence learning, VAS – Visual analogue scale.

### 2.3. Motor Sequence Learning (MSL) Task

The MSL Task was programmed using Psychopy (Open Science Tools Ltd, Nottingham) [32] and utilised a joystick (Thrustmaster T.16000M FCS Flightstick) in the dominant hand to move a cursor between eight different targets positioned in one of four locations on a computer screen (Figure 2), visiting each location twice. Evidence of face validity of this task has been described by Khatibi et al. (2022). The authors demonstrated increased task performance and learning related changes in neural activity in brain and spinal cord, across a period of learning. The trial started when participants initially hovered the cursor over a small blue central target for 0.5 seconds which reflected the central position of the joystick. The central target would then disappear, and the first of the eight targets would appear.

**Fig 2:**
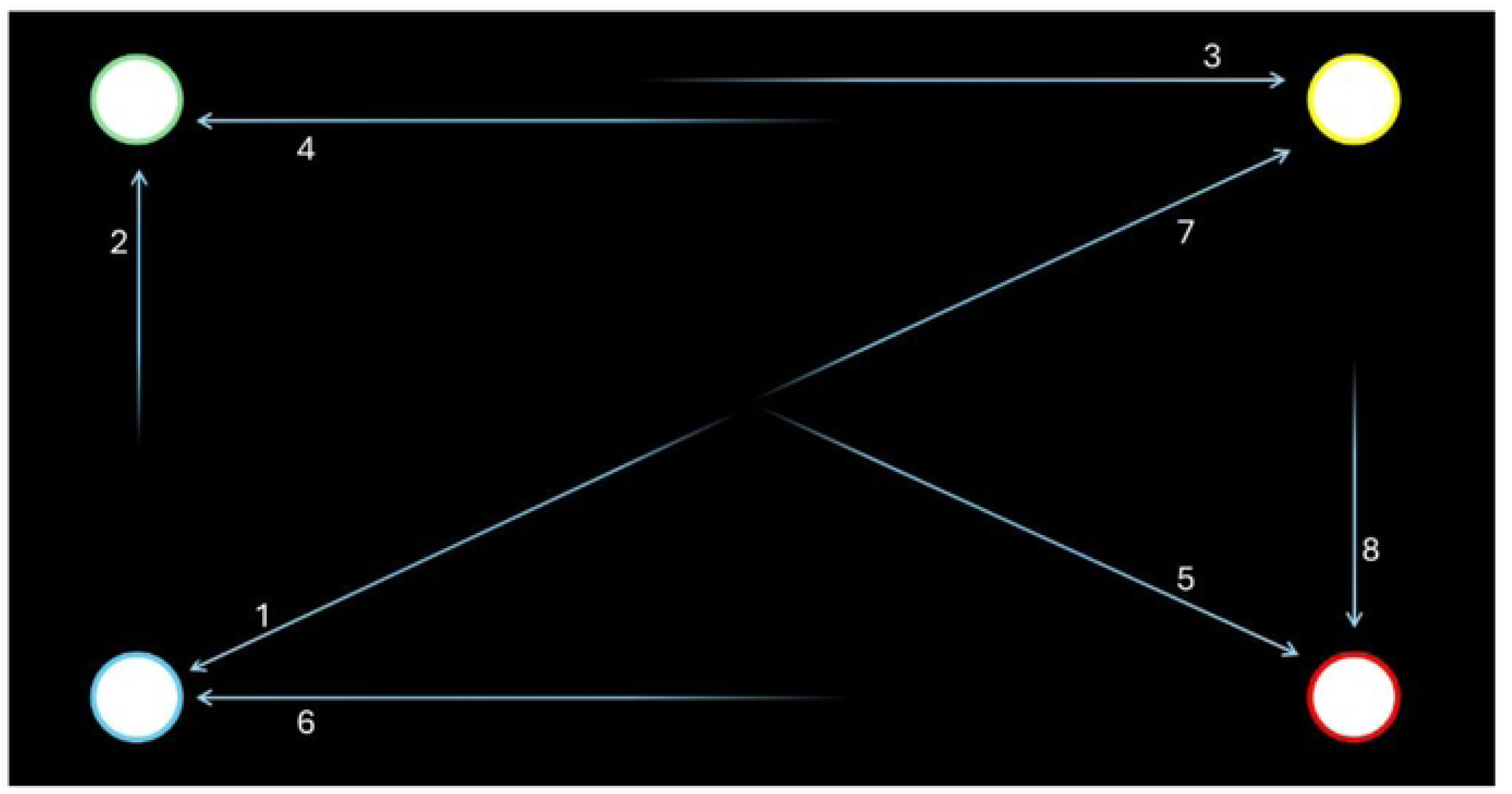
Position and orders of targets. The arrows and numbers demonstrate the order of the learnt sequence.

The participant would then move the cursor to the new target and was required to sustain that position for 0.5 seconds, to avoid participants ‘shooting’ through the targets. That target would subsequently disappear and trigger the next target to appear. This would continue until all eight targets were visited. All participants completed 150 sequences across 10 trials which is consistent with the duration of training used in previous motor learning research [33, 34]. The participants received a two-minute break between trials. Finally, in an effort to maintain motivation during the task an auditory beep sounded if the participant completed the sequence faster than a pre-selected time. The pre-selected time was calculated from pilot data and was based on the lowest 33% percentile (fastest) of all sequences completed. No other feedback was provided.

Prior to completing the MSL session participants were given time to read written instructions outlining the task and completed five random sequences during a familiarity session. The written instructions stated that participants must ‘complete the task as quickly and accurately as they can’. This statement was repeated at the start of each trial.

### 2.4. Experimental movement-contingent pain paradigm

Electrical stimuli was administered via two self-adhesive electrodes (2.5 × 2.5 cm, CDES003545, Spesmedica, Genova, Italy) to the dorsal surface of the participants right wrist using a constant-current stimulator (Digitimer DS5 Isolated Bipolar Constant Current Stimulator, Welwyn Garden City, Hertfordshire, UK) controlled by an analogue signal generated by a NI multifunction data acquisition board (PCI-6229, 16 bit resolution, 2000Hz, National Instruments, Austin, TX, USA). Prior to the MSL paradigm, individual stimulus intensities were established for each of the participants using an increasing stimulus paradigm adapted from Meulders et al (2011). An increasing stimulus was applied in 0.5 increments. Participants were asked to rate the sensations caused by the stimulus on a scale of 0-10. (‘‘0’’ indicating that they felt no pain, ‘‘1’’ indicating that the sensation was starting to become aversive and slightly painful (pain threshold), and ‘‘10’’ indicating the worst pain that they could imagine). Intensities scored just below one out of ten was utilised for the non-noxious control group and intensities inducing scores of five out of ten were used in the intervention groups. Previous literature has used a target score of five out of ten in experimental pain groups as it is deemed to be a ‘moderate pain that requires effort to tolerate [36] and therefore has the potential to require allocation of attentional resources. Once a score of six out of ten was reported the stimulation was returned to zero and the process was completed twice more. An average of the three intensities was calculated and used during the experiment.

The electrical cutaneous movement-contingent experimental pain paradigm was adapted from Gallina et al (2021). An electrical stimulus which increased in intensity based on the joystick movements was control using MATLAB (MathsWorks, USA). Once the participant moved beyond zero, in a defined direction on the y axis the stimulation would start, increasing proportionally in relation to its position on the y axis from zero, peaking on reaching a target and disappeared completely once the cursor returned to zero on the y axis. This setup was chosen to give the participants the feeling that the pain was evoked by the movement of the wrist. The defined direction was randomized for participants, within each group, between up or down movements of the joystick. The perceived ‘level of pain’ during each trial was collected using a visual analogue scale (VAS) following the trial.

### 2.5. Eye Tracking

Eye kinematic data was collected using Eyelink 1000 plus (SR Research Ltd) using a sample rate of 500hz. The Eyelink 1000 hardware was controlled by a ‘host’ computer which was linked to a ‘display’ computer which ran the MSL task enabling Psychopy to send messages to the host computer to timestamp when targets appeared during the task. The tower-mount setup (Figure 3) was utilized to maintain head position and standardize the angle of the head in relation to the screen, throughout the trials.

**Fig 3:**
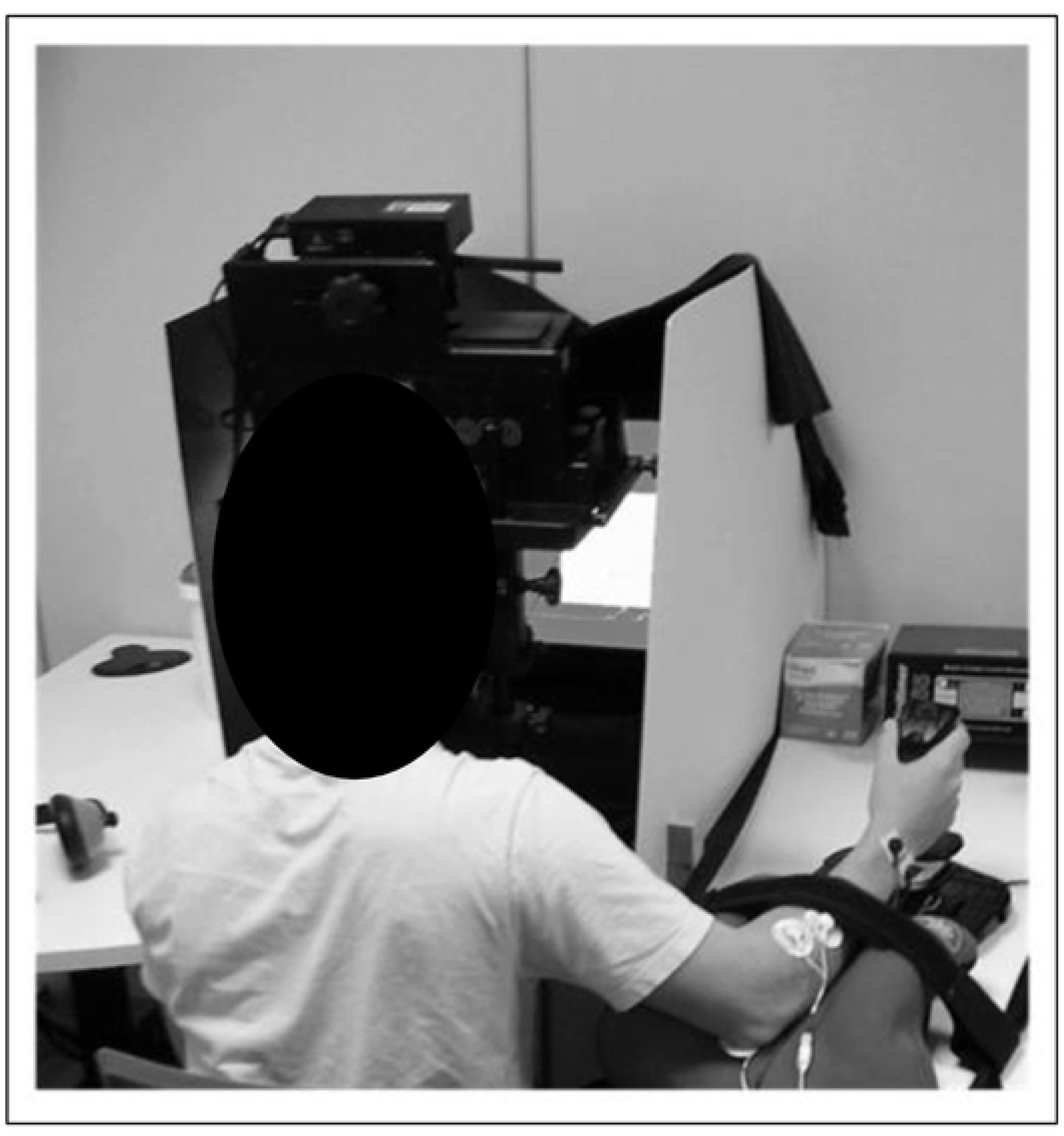
Setup of eye tracking.

Changes in accuracy over time is known as drift and will impact on the quality of data collection. To minimise drift, calibration and validation of the Eyelink 1000 was completed prior to each trial. Settings for the Eyelink 1000 and calibration and validation procedures can be seen in (S3 Table). Validation was accepted if average accuracy of all nine points was less than 0.5 degrees and no single point was more than one degree of error [37]. Immediately after the validation was complete the motor task was started. Participants were asked to keep their heads still from the start of the calibration process to the end of the trial.

### 2.6. Outcome Measures

#### 2.6.1. Task performance measures

Changes in behaviour across training was measured using the following temporal and spatial measures.

1. Speed – defined as time to complete sequence or time to reach targets.
2. Accuracy – defined as the average distance points (coordinates) were away from a polygon representing the shortest and most direct route from one target to the next.
3. Dimensionless squared jerk 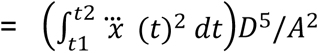

*A = Overall distance to target D = t_2_-t_1_ (Overall duration to target)*

#### 2.6.2. Eye gaze measures

The eye gaze indices were selected to provide insights into the different attentional mechanism. Outcome measures selected for this study are outlined in Table 1 and have been used previously in studies using eye gaze technology [38].

**Table 1:** Eye gaze indices analysed in this study.

| Attentional mechanisms/processes | Outcome measure | Definitions |
| --- | --- | --- |
| Initial orientating | Target entry time | The time difference between the target appearing and the eye movement first fixating on the target. |
| Attentional engagement | Fixation count | The number of fixations on individual targets per sequence. |
| Attentional maintenance | Dwell time | The overall dwell time on individual targets per sequence. |
|  | First fixation duration | The duration of the first fixation on targets following the target appearing. |
| Difficulty disengagement | Target exit times | The time difference between the target disappearing and the eye movement ending the last fixation on the target. |

Improvements in efficiency of attentional control would be indicated by a reduction in each of the above measures between early and late training.

### 2.7. Data Processing

#### 2.7.1. Processing of spatial and temporal task performance data

Temporal and Spatial data was collected via Psychopy software (Open Science Tools Ltd, Nottingham) and MATLAB using SIMULINK (MathsWorks, USA), respectively. Data from each trial was visually inspected and was then segmented into individual sequences and individual targets using bespoke MATLAB (MathsWorks, USA) scripts. Data for the time taken to reach targets was normalised with respect to the average time for all participants for diagonal, vertical and horizontal transitions. To calculate accuracy, the distance of each point from a polygon representing the shortest and most direct route was calculated (S1 Fig) and then normalised in relation to time by dividing by the number of points in a single transition. When calculating jerk, the decision was made to down sample joystick coordinate data from 500hz to 30hz to minimise the impact of multiple sustained periods of arrest on the measurement of jerk. Following down-sampling, coordinate data was analysed using bespoke MATLAB (MathsWorks, USA) scripts resulting in calculations of Euclidean displacement (d), speed (v) and acceleration (a) which was inputted into the equation above.

#### 2.7.2. Processing eye gaze data

The EyeLink Data Viewer software (SR research, version 4.2.1) was used for the following data processing. During data processing the data was visually inspected and discarded if either of the following was true.

1. Gaze moved outside the parameters of the screen during any part of the trial.
2. More than 15% of the trial data was missing across each trial.

Segregation of eye gaze data into the individual transition times relating to each target was achieved using the time stamps sent from Psychopy. Regions of Interest (ROIs) were defined for each target location (Figure 4). A fixation was defined as a gaze that remains stable for a duration of more than 100ms.

**Fig 4:**
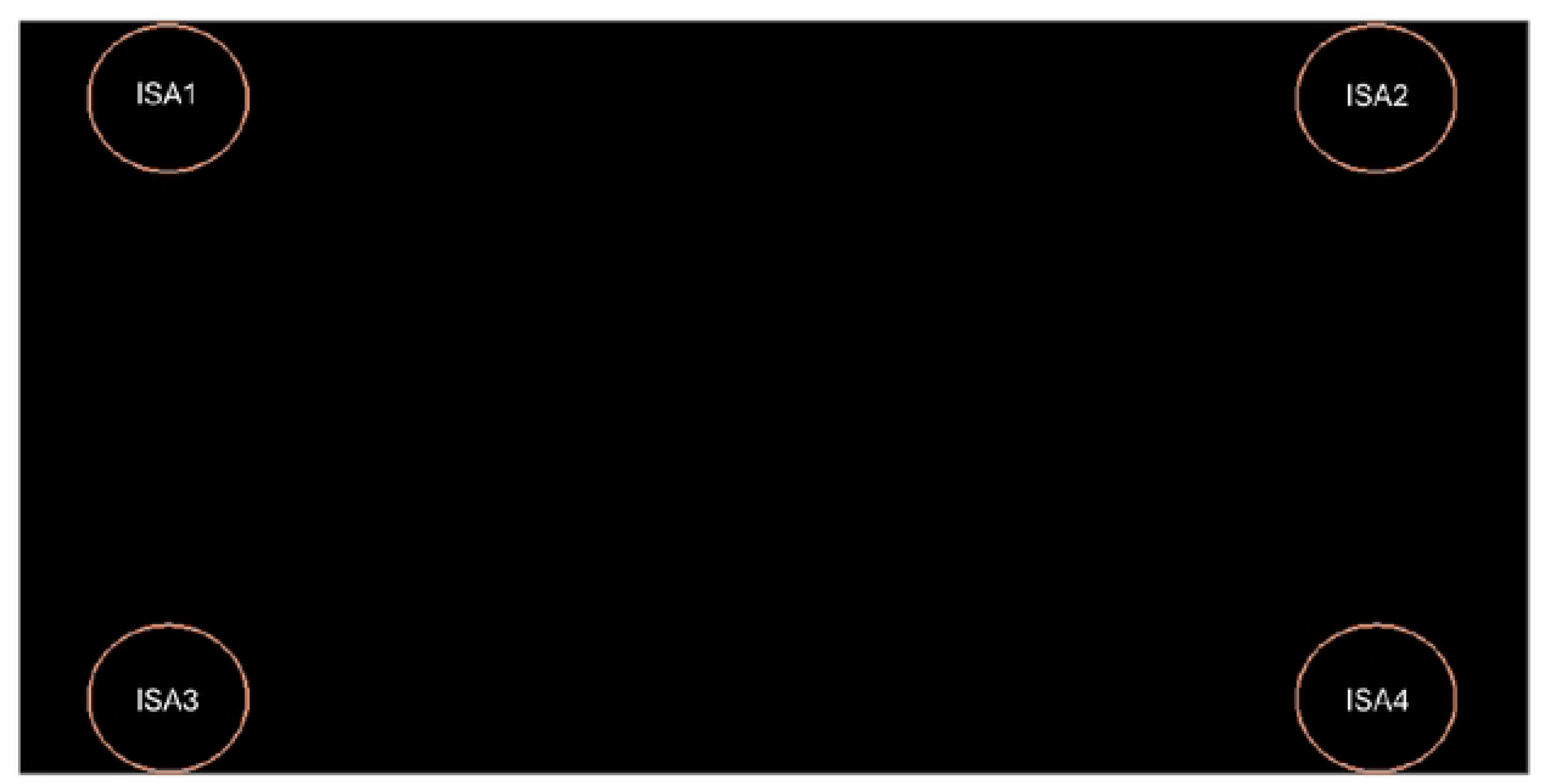
Diagram representing location and size of Regions of interest.

For all task performance outcome measures, early (first and second trials), mid (fifth and sixth trials) and late (ninth and tenth trials) training epochs were identified for each participant. Due to the quality of eye gaze measures, only the early (first and second trials) and late (ninth and tenth trials) training epochs were calculated for analysis. The first block of the first trial for each participant was removed to avoid the possible impact of task familiarisation. Averages were calculated for each epoch so comparisons could be made across training.

### 2.8. Statistical Analysis

Baseline demographics and self-reported measures were collated and following analysis for data normality, one-way ANOVAs were used to explore differences between groups. Pain scores across trails did not meet requirements for normality, so to explore the robustness and face validity of the movement-contingent pain paradigm the following analysis was completed. To establish whether the electrical stimulus was able to induce a sensory experience similar to the target sensation, the mean VAS scores for ‘level of pain’ for each trial for each group was compared to the target value using the Mann-Whitney U test. Following this, the consistency of the induced sensory experience was assessed across time for each group using the Friedman test. Differences in machine stimulus intensity was compared using the Wilcoxon test to confirm that different intensities were used for intervention and control groups. Finally, a Kruskal-Wallis H Test was completed to assess the difference between groups at each time point for the ‘level of pain’.

To explore the impact of within-subject variables (epoch and targets) and between subject variables (group allocation) a mixed repeated ANOVA was applied to log transformed data for temporal and spatial performance measures and eye gaze indices; fixation count, first fixation duration and target entry times. A mixed repeated ANOVA was also used to explore the impact of stimulation direction on the above outcomes. Dwell time and target exit times violated the above parametric test requirements and therefore the Wilcoxon rank test was used to assess impact of within subject factors and Mann U Whitney was utilised to explore between subject variables. Spearman’s rank correlation coefficient was used to explore the relationships between eye gaze measures and task performance measures, self-reported questionnaires and demographic data.

## 3. Results

### 3.1. Demographics

Age (F(2, 63) = .764, *p* = .470), stress (F(2, 63) = .582, *p* = .562), depression (F(2, 63) = .2.112, *p* = .130), anxiety (F(2, 63) = 1.584, *p* = .213) and fear avoidance (F(2, 63) = .004, *p* = .996) were similar for all groups. The pain-intervention group reported significantly higher levels of pain catastrophizing (F=(2, 63) = 6.007, *p* = .004) compared to the control and intervention group (Table 2).

**Table 2:** Mean and range for demographic data and self-reported measures for all three groups.

|  |  | Control<br>(n=28) |  |  | Intervention<br>(N=24) |  |  | Pain-intervention<br>(n=14) |  |  |
| --- | --- | --- | --- | --- | --- | --- | --- | --- | --- | --- |
|  |  | Mean | Range |  | Mean | Range |  | Mean | Range |  |
|  |  |  | Min | Max |  | Min | Max |  | Min | Max |
| Age |  | 24.68 | 18 | 31 | 24.25 | 19 | 34 | 25.93 | 19 | 41 |
| Gender (F:M) |  | 13:15 |  |  | 12:12 |  |  | 6:8 |  |  |
| Self-reported questionnaires |  |  |  |  |  |  |  |  |  |  |
| Pain catastrophizing |  | 10.46 | 0 | 43 | 9.57 | 0 | 22 | 20.07 | 0 | 45 |
| DASS | Stress | 4.68 | 0 | 17 | 3.61 | 0 | 11 | 4.21 | 0 | 9 |
|  | Anxiety | 2.64 | 0 | 7 | 1.61 | 0 | 8 | 2.64 | 0 | 6 |
|  | Depression | 2.64 | 0 | 17 | 1.39 | 0 | 8 | 3.21 | 0 | 8 |
| Fear Avoidance |  | 74.86 | 30 | 121 | 74.43 | 41 | 99 | 74.36 | 32 | 117 |
| Attentional control – Total |  | 41.83 | 24 | 58 | 39.19 | 28 | 49 | 39.80 | 23 | 51 |
| Attentional control – Shifting |  | 24.09 | 17 | 34 | 23.56 | 15 | 32 | 24.20 | 11 | 32 |
| Attentional control – Focusing |  | 17.74 | 6 | 25 | 15.63 | 5 | 23 | 15.60 | 8 | 23 |

### 3.2. Movement Evoked Pain Paradigm

‘Level of pain’ for the intervention group were significantly lower for all trials than the target score of five indicating a possible deviation from the planned intensity of the pain experience for this group. In contrast, there was no significant difference between the ‘level of pain’ reported by the pain-intervention group and the target score across all trials (Figure 5). Friedman test demonstrates that there was no significant change across time for ‘level of pain’ for control (*p* = .087), intervention(*p* = .554) or pain-intervention (*p* = .335) groups.

**Fig 5:**
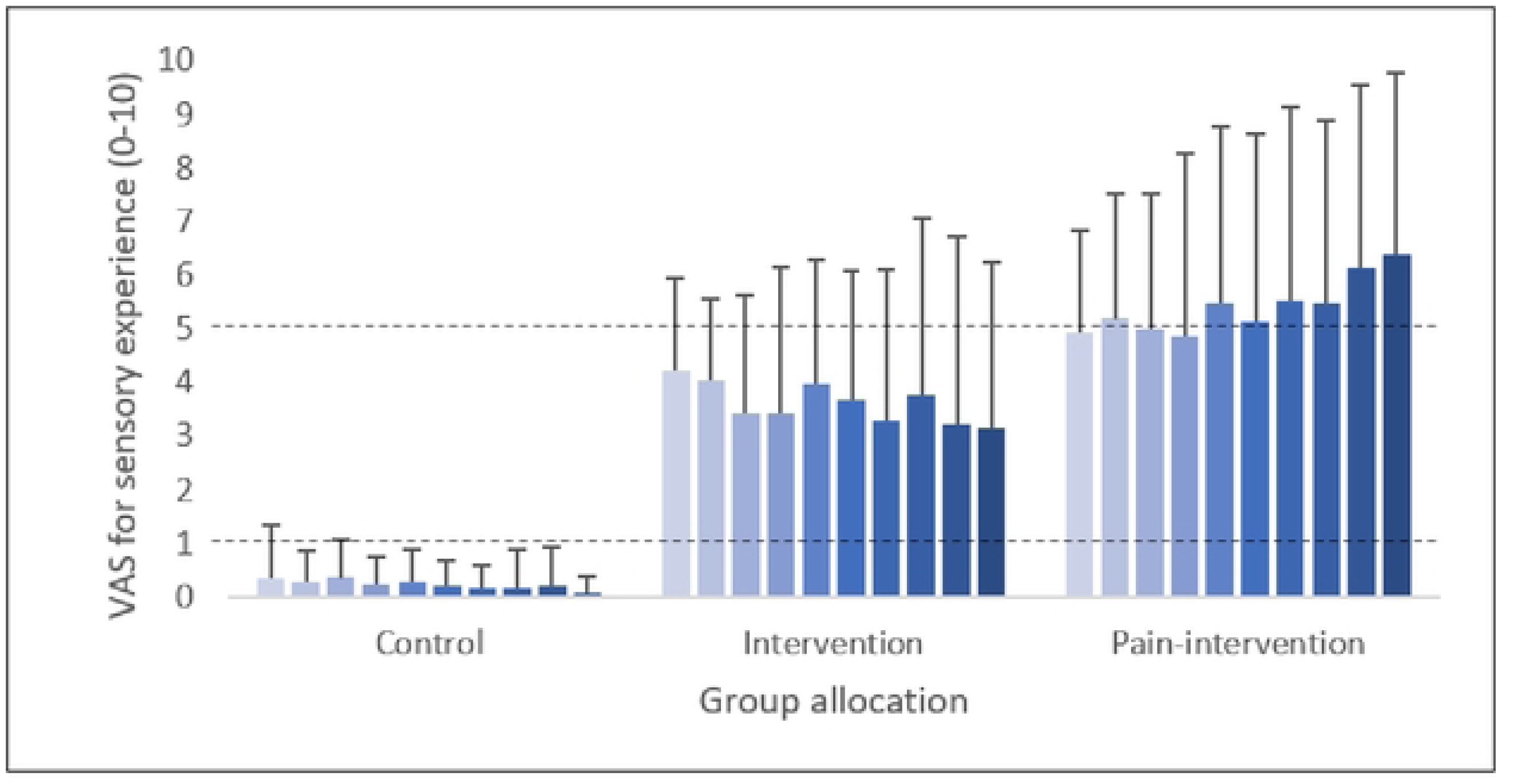
Bar charts demonstrating median and interquartile range of VAS scores for ‘level of pain’ for each trial. Target values of 1/10 and 5/10 have been marked on the ‘level of pain’ bar chart.

The Kruskal-Wallis H Test showed there was a significant difference in VAS for the ‘level of pain’ reported during all trials (*p* < .001) between the control, and the intervention and pain-intervention. When pooling all groups, stimulus intensities to induce target level sensations of 1/10 and 5/10, established during the increasing stimulus protocol, were significantly different (Wilcoxon signed-rank test; *N* = 72; *z* = 7.375, *p* < .001). The above analysis confirms the sensory experience reported by all groups was consistent across the ten trials and that the control group was exposed to a different condition compared to the two intervention groups. This provides support for the use of the movement evoked pain paradigm used in this study as a valid method for inducing stable experimental pain. Although it must be noted that the intensity of the painful sensation in the intervention group was lower than set out in the methodology.

### 3.3. Task Performance measures

#### 3.3.1. Time to Complete Sequence

When comparing time to complete sequence (Figure 6) between early, mid and late training epochs a significant main effect of time was observed (F(1, 63) = 274.292, *p* < .001, n_p_^2^ = 0.813) indicating motor performance improved across time for all groups. The significant improvement in time was observed both between early and mid-training and mid and late training. Comparison across groups showed no main effect of group allocation on performance (F(2, 63) = 2.469, *p* = .093) and no time by group interaction (F(1, 63) = 0.139, *p* < .955). This indicates all groups ‘time to complete sequences’ improved over time and this was not impacted by the experimental pain regardless of the presence of a history of pain.

**Fig 6:**
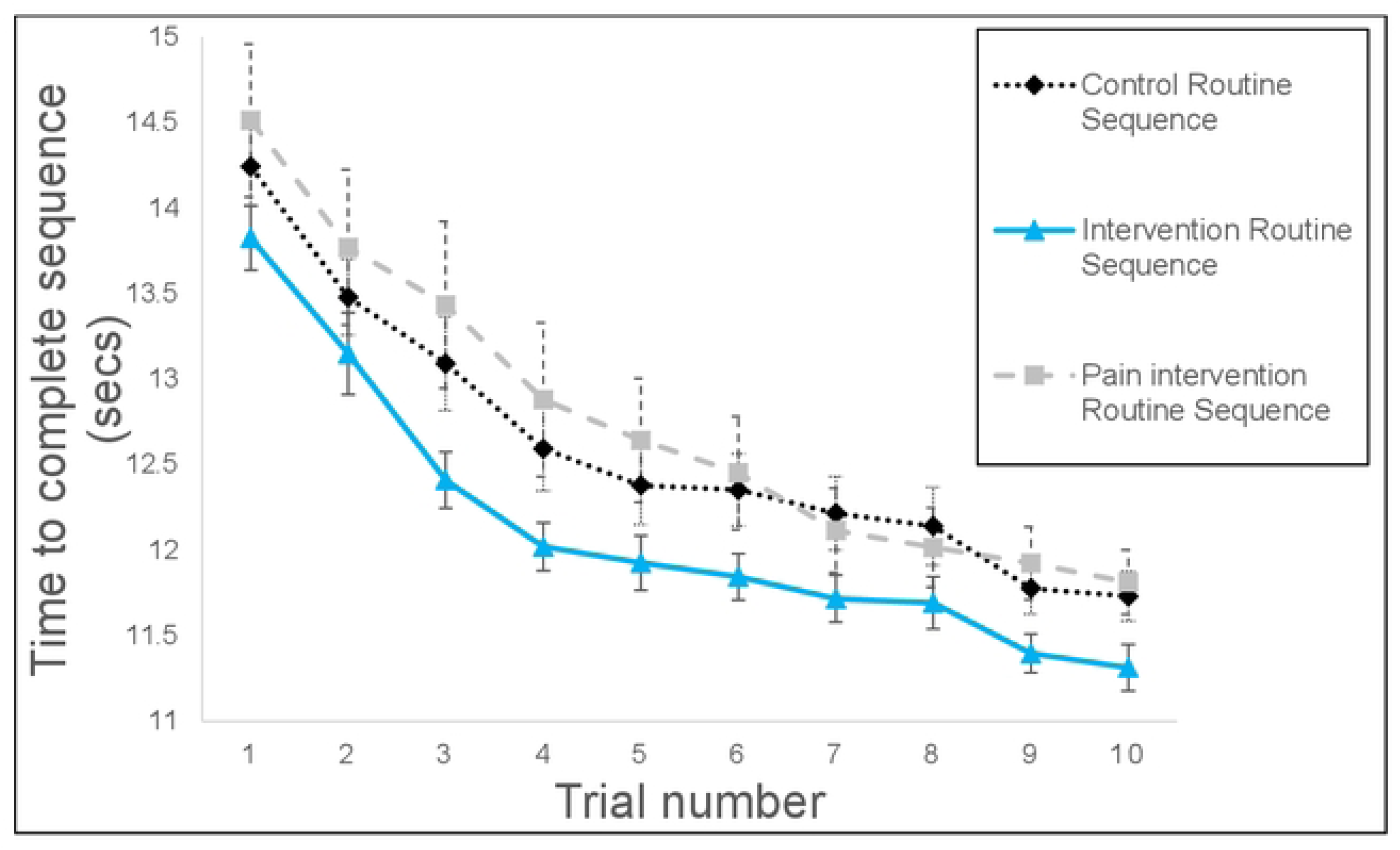
Mean and standard error for change in ‘time to complete sequence’ across the 10 trials for all three groups.

#### 3.3.2. Accuracy

There was a main effect of time (F(1, 63) = 9.125, *p* < .001, n_p_^2^ = 0.127) on *accuracy* indicating improvements across time in all groups, although the effect size was small (Figure 7a). In contrast to the temporal data, improvements were only seen between mid and late training. A main effect of group was observed on accuracy (F(1, 63) = 11.634, *p* < .001, n_p_^2^ = 0.270). The intervention group was more accurate than the control or pain-intervention group. An interaction between condition and time failed to demonstrate significance (F(1, 63) = 2.363, *p* < .057) suggesting that the intervention group was more accurate at all time points during the training and the group effect did not represent superior motor learning across the training.

**Fig 7:**
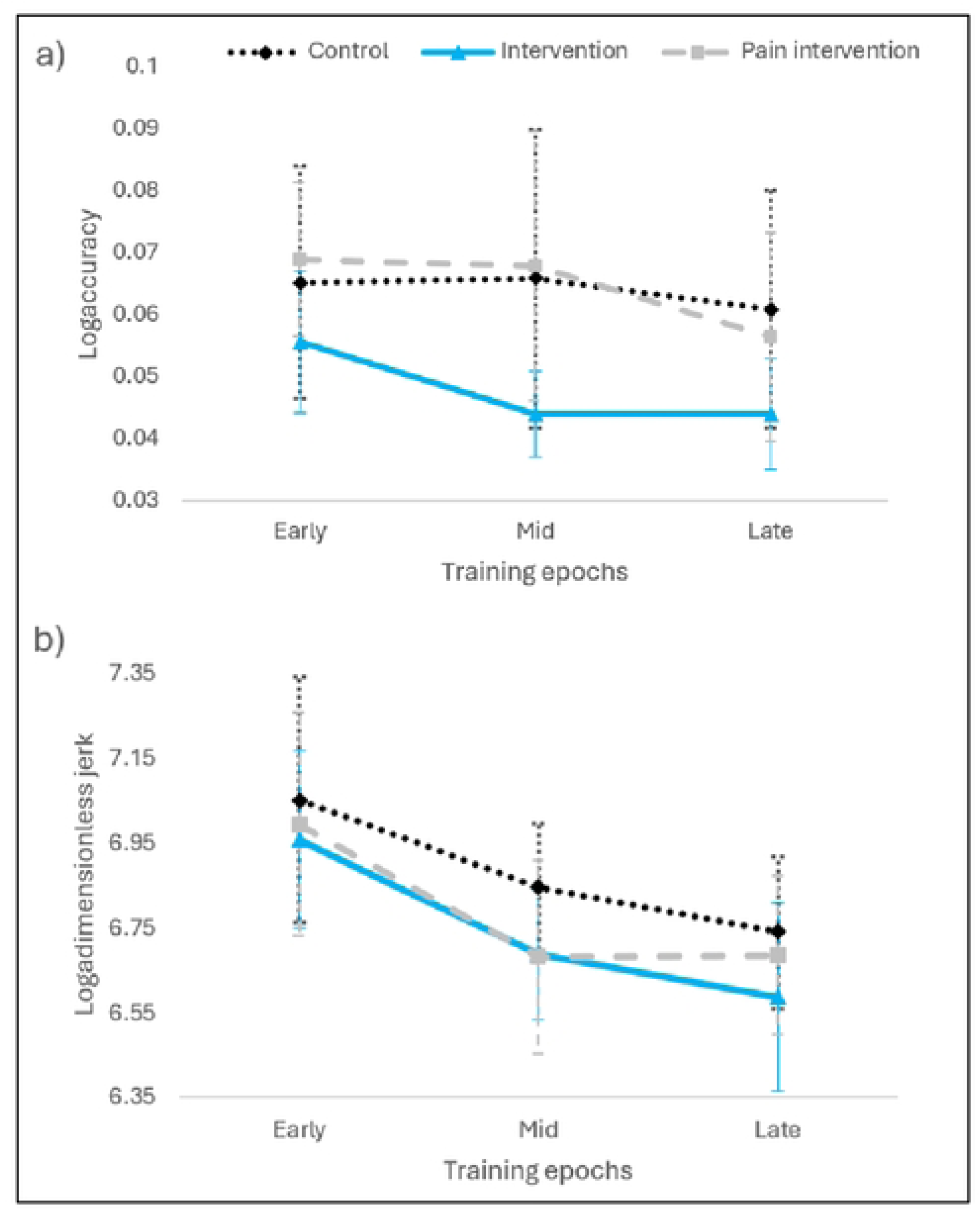
Mean and standard deviation for a) Accuracy and b) Dimensionless jerk across training for all three groups.

#### 3.3.3. Dimensionless Jerk

There was a significant main effect of time on dimensionless jerk for whole sequence (F(1, 63) = 57.723, *p* < .001, n_p_^2^ = 0.486)(Figure 7b) suggesting there was an improvement in smoothness of movement across training for the whole sequence irrespective of group. There was also a main effect of group for whole sequence (F(1, 63) = 4.268, *p* < .018, n_p_^2^ = 0.123) suggesting an impact of experimental pain on smoothness of movement. There was no interaction between time and group for whole sequence (F(1, 63) = 0.774, *p* < .544), indicating there was no difference in change in smoothness over time between the groups. The results suggested experimental pain increased smoothness in the intervention group when compared to the control group only.

#### 3.3.4. Participant Task Engagement

To explore whether task performance during the task was impacted by whether the target was associated with experimental pain or not, a further subgroup analysis was performed on all measures for individual targets. No effect of stimulus direction was observed for ‘time to reach targets’ (Control: F(1, 26) = 0.660, *p* = .424, Intervention: F(1, 22) = 0.440, *p* = .514, Pain-intervention: F(1, 14) = 0.001, *p* = .974), accuracy (Control: F(1, 26) = 0.015, *p* = .903, Intervention: F(1, 22) = 1.106, *p* = .304, Pain-intervention: F(1, 14) = 0.106, *p* = .750) or dimensionless jerk (Control: F(1, 26) = 1.422, *p* = .244, Intervention: F(1, 22) = .560, *p* = .462, Pain-intervention: F(1, 14) = 0.085, *p* = .776) for individual targets in all groups. This provides some indication that pain did not impact on participants engagement in the task.

### 3.4. Attentional Control

#### 3.4.1. Attention across learning

Figures 8-10 show the changes in eye gaze measures across time for each of the eight targets for each group. To aid interpretation the following section will be divided into the attention mechanisms presented in Table 1.

**Fig 8:**
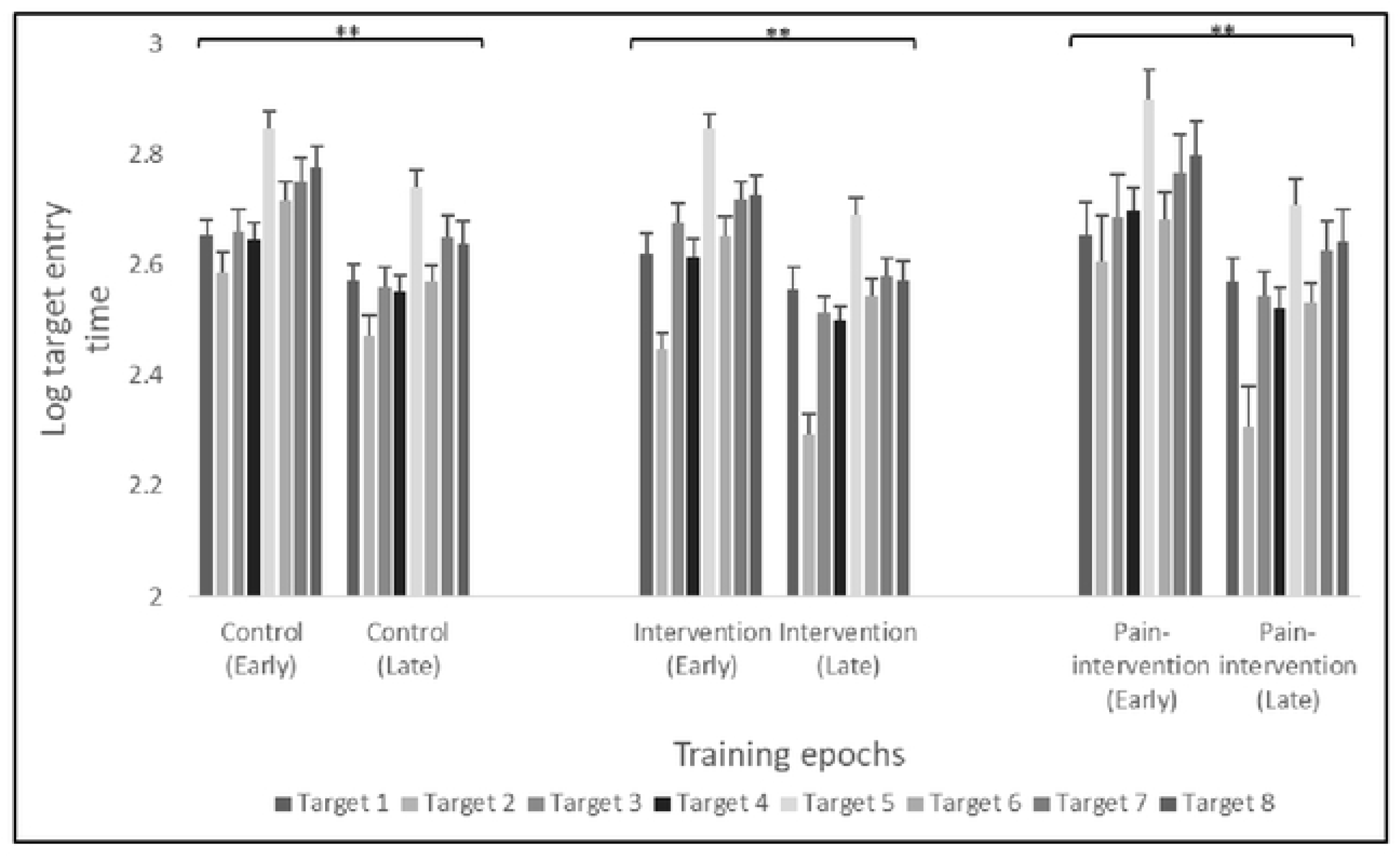
Mean and standard error for target entry time for all 8 targets for Control, Intervention and Pain-intervention groups. * p< 0.05, ** p < 0.01.

**Fig 9:**
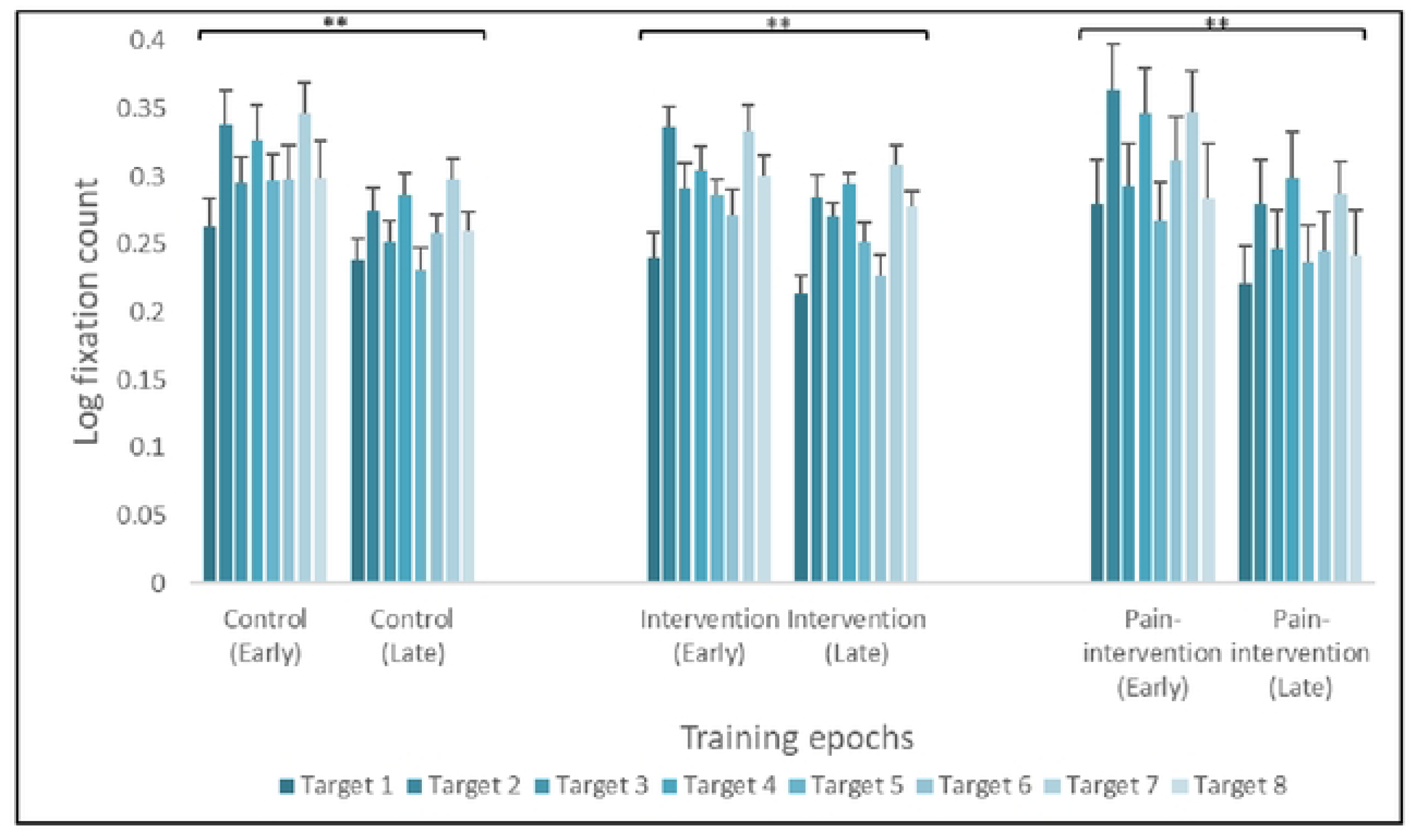
Mean and standard error for fixation count for all 8 targets for Control, Intervention and Pain-intervention groups. * p< 0.05, ** p < 0.01.

**Fig 10:**
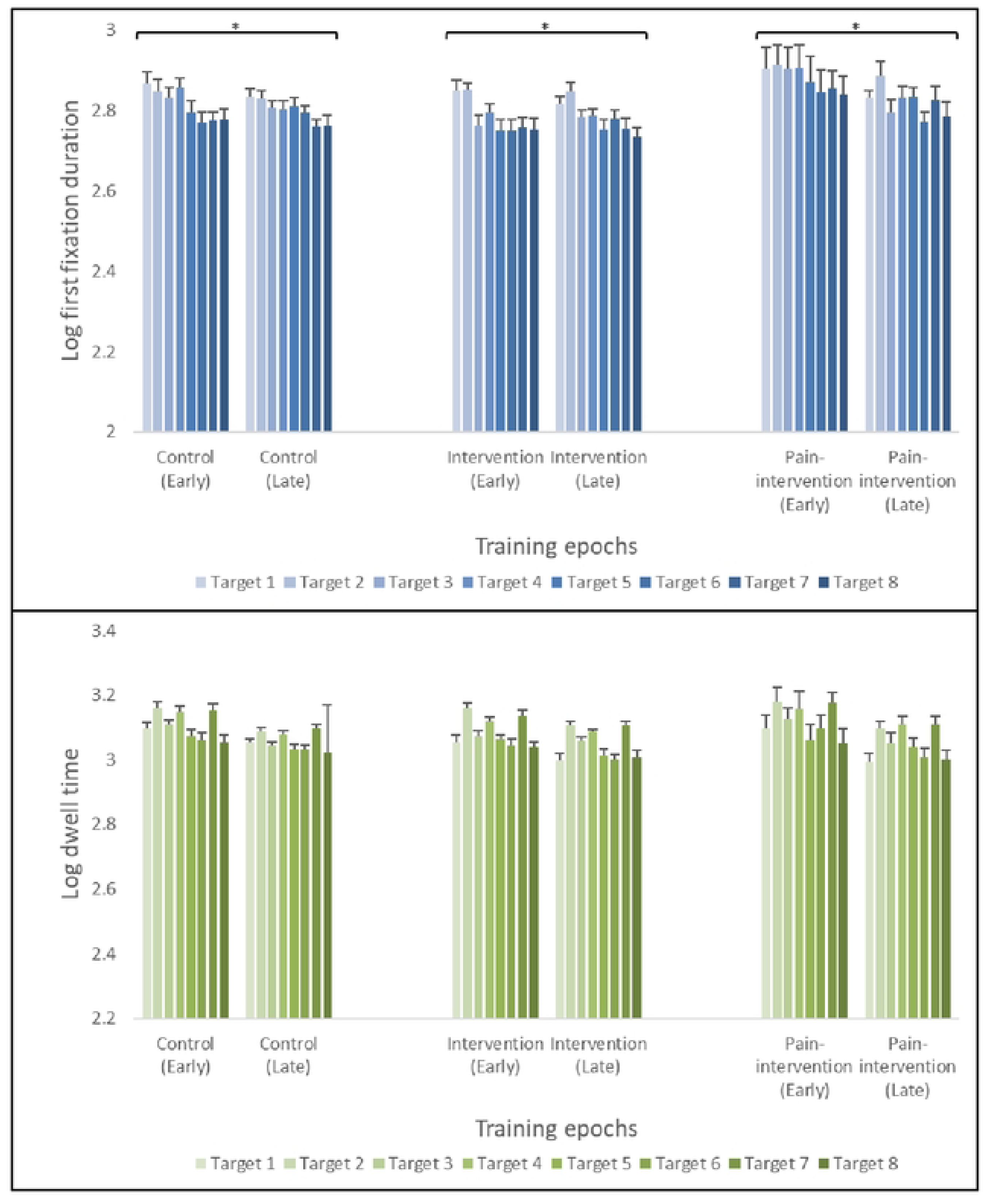
Mean and standard error for first fixation duration and dwell time for all 8 targets for Control, Intervention and Pain-intervention groups. * p< 0.05, ** p < 0.01.

##### Initial Orientating

A significant main effect of time was observed for target entry time (f(1, 46) = 116.607, *p* < .001, n_p_^2^ = 0.713). The observed significant reduction in the time taken to enter targets suggests an improvement in efficiency of initial orientating of attention for all targets in response to task stimuli across training. A significant main effect of target was also observed (f(1, 46) = 53.624, *p* < .001, n_p_^2^ = 0.533) suggesting target entry time was dependent on both time visited and location of the target. There was no observed effect of sequence order on the sequence on target entry time. A significant moderate positive correlation was observed between target entry time and time to target during early training for both the control group (rs= .625, *p* = .001) and pain-intervention group (rs= .661, *p* = .038) but not the intervention group (rs= .103, *p* = .694). This provides some evidence that the process of initial orientating of attention may work in parallel with motor mechanisms during early acquisition of motor skills, but not in all populations and not under all experimental conditions.

##### Attentional Engagement

A significant main effect of time was observed for fixation count (f(1, 46) = 19.505, *p* < .001, n_p_^2^ = 0.293). A reduction in number of fixations was observed during late training compared to early training for all targets and all groups. Similar to initial orientating, a significant main effect of target was also observed on attentional engagement (f(1, 46) = 20.335, *p* < .001, n_p_^2^ = 0.302). Fixation count was not impacted by where the target appeared in the sequence. Fixation count showed a significant moderate positive correlation with time to target at both early and late training for the control (early, rs= .459, *p* = .027, late, rs= .576, *p* = .004) and intervention (early, rs= .659, *p* = .004. late, rs= .566, *p* = .018) groups. This indicates increased efficiency of movement was associated with a smaller number of fixations on the target throughout training. This transition from higher to lower numbers of fixations may be a characteristic of increased attentional efficiency. Similar to initial orientating this relationship may vary depending on the population as no significant correlation was observed in the pain-intervention group (early, *p* = .960. late, *p* = .556).

##### Attentional Maintenance

A significant main effect of time was observed for first fixation duration (f(1, 46) = 4.788, *p* < .034, n_p_^2^ = 0.092) but not for dwell time (f(1, 46) = 1.966, *p* < .167) indicating fixation duration may prove to be a more appropriate eye gaze measurement to explore attentional maintenance across training than dwell time. Interestingly, early training dwell time, not first fixation duration, had a significant moderate positive correlation with time to target in the intervention group (rs= .696, *p* = .002) and a significant moderate positive correlation with accuracy in the pain-intervention group (rs= .733, *p* = .016). A significant main effect of targets was observed for both first fixation duration (f(1, 46) = 12.152, *p* < .001, n_p_^2^ = 0.205) and dwell time (f(1, 46) = 41.573, *p* < .001, n_p_^2^ = 0.469) but no impact of sequence order was observed.

##### Disengagement

Data on target exit times and therefore disengagement from stimuli across training provided non-significant results.

#### 3.4.2. Impact of Experimental Pain on Attentional Control

No significant main effect of condition on eye gaze measures of attention control was observed (fixation count, f(1, 46) = .024, *p* = .977, first fixation, f(1, 46) = 2.460, *p* = .096, dwell time, f(1, 46) = .011, *p* = .989, target entry time, f(1, 46) = .977, *p* = .384). No significant difference between groups at any time point was observed for target exit times (p-value ranged from .247 to .815). An interesting finding in the intervention group was, total pain catastrophizing and the associated three sub categories all had a moderate positive correlation with fixation count during early training (Total, rs= .699, *p* = .003, Ruminating, rs= .579, *p* = .019, magnification, rs= .617, *p* = .011 and helplessness, rs= .686, *p* = .003) but total fear avoidance (rs= - .517, *p* = .040) and severe sub category of fear avoidance (rs= -.518, *p* = .018) had a moderate negative correlation with fixation count during late training (Figure 11). Therefore, participants in the healthy experimental pain group, who reported higher levels of pain catastrophizing had a higher number of fixations during early training on all targets. Conversely, in the same group, higher levels of fear avoidance resulted in a lower number of fixations during later stages of learning. No correlations were present between self-reported measures and eye gaze measures for initial orientating, attentional maintenance or disengagement for any group.

**Fig 11:**
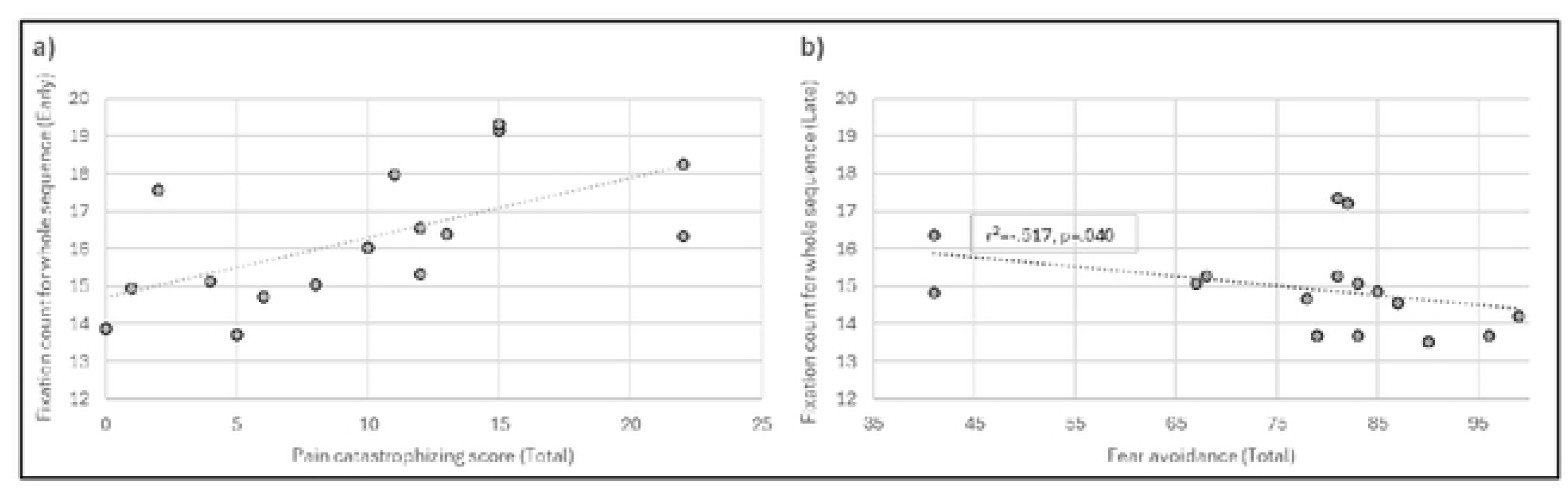
Scatter graphs representing correlation between fixation count and 1) pain catastrophizing, 2) fear avoidance in the intervention group.

## 4. Discussion

The aim of this study was to investigate whether an electro-cutaneous movement-contingent experimental pain paradigm can create a consistent pain experience across multiple motor learning trials and explore the impact of this pain paradigm on task performance improvements across motor learning and on the pattern of attentional allocation, measured using gaze indices.

The results of this study demonstrate that the novel electro-cutaneous stimulation paradigm was able to induce a stable pain experience across multiple trials which clearly differed from the non-noxious electro-cutaneous control. This was despite the level of pain reported in one of the intervention groups differing from the target level. Reflecting findings with tonic experimental pain paradigms [18], movement contingent experimental pain did not impact on improvements in task performance across motor training, but did result in improved accuracy and increased smoothness of movement at all time points in healthy participants. Three out of the five eye gaze indices measured in this study demonstrated significant changes across motor learning. Initial orientating, attentional engagement and attentional maintenance, showed significant differences from early training to late training. All measures indicated a transition to a more efficient allocation of attention. This included a decrease in the time it takes for gaze to initially fix on a target, a decrease in the duration of the first fixation on a target and a decreased number of times gaze fixes on a target. Movement-contingent experimental pain does not impact on changes in attentional control across a period of MSL. The above finding was consistent regardless of the attentional process under investigation or whether the population experienced persistent pain.

### Movement Contingent Pain

Electro-cutaneous stimulation paradigms have been plagued with issues around habituation [22, 39] and lack of specificity of activation of neuronal afferents [17]. These concerns have led to questions on whether electro-cutaneous stimulation can induce a valid and reliable pain experience. Gallina et al, (2021) proposed the use of low frequency sinusoidal waves to overcome these issues and demonstrated no habituation over 60 seconds using this approach. The findings of this study extend this to demonstrate a stable pain experience across ten trials of motor learning, suggesting this approach could be viable to explore the impact of pain experience on prolonged activity. The ability of electro-cutaneous stimulation to induce a pain intensity similar to the target level was best demonstrated for the participants with persistent pain. Exposure to electro-cutaneous stimulation in healthy participants resulted in a stable ‘level of pain’ across the ten trials but at a significantly lower level than the target score set in the methodology. The healthy intervention group reported low levels of pain catastrophizing, and this maybe reflected by the lower levels of pain being reported. Further research is needed collecting data on pain levels at more regular intervals to provide more detailed information on the timeline of pain experience during motor training.

The range of ‘levels of pain’ reported across both intervention groups for all trials was consistent with pain levels reported by previous research using both capsaicin and hypertonic saline injections to explore the impact of pain on motor learning [9–11, 13, 40–42]. This provides some support that the pain levels induced with electro-cutaneous stimulation is comparable to that induced using other common experimental pain paradigms with the added benefit of being contingent on movement.

### Pain Interference with Motor Learning

This study found that participants exposed to movement–contingent experimental pain, regardless of whether they reported a history of persistent pain or not, demonstrated improvements in ‘time to complete sequence’ comparable to improvements seen in participants experiencing non-noxious electro-cutaneous stimulation, during a session of MSL. These findings are in line with previous research demonstrating no effect of pain on improvements in temporal measures of task performance across learning [9, 10, 13, 42]. The findings of the present study suggest that movement-contingent experimental pain does not impact on improvements in accuracy across a session of MSL. This is consistent with findings from the majority of previous research exploring the impact of pain on accuracy measures of task performance during acquisition [18]. Only two previous studies have reported an impact of experimental pain on motor learning during acquisition [10, 40]. In contrast to the present study, Dancey et al. (2016a) utilised a simple typing task; the lack of improvement of performance observed in the group experiencing pain could be partly down to the high accuracy scores at baseline and the resultant ceiling effect for this measure. Studies minimising the risk of ceiling effects by utilising more complex motor learning tasks [11, 13, 43], like the one used in the present study, have more than often failed to find the same impacts on learning. Boudreau et al. (2007) is the only study to report reduced improvements in accuracy in the presence of experimental tonic pain during a complex tongue protrusion task. It has been argued that the process of application of capsaicin by Boudreau and colleagues had the potential to impact engagement in the task and that disengagement from the task may have explained the reduction in learning. No attempt was made to monitor or adjust for this and therefore caution needs to be taken with interpreting their results. In the present study, accuracy scores on reaching individual targets were similar regardless of whether the target was associated with pain or not, suggesting pain did not directly influence engagement in the task. This provides some evidence that if all participants engage in the task, similar levels of motor learning are observed despite the presence of pain.

Increased accuracy in the presence of experimental pain at all time points of motor learning, as seen in the intervention group in this study, has been reported by previous research [11, 13, 43]. Hypotheses suggest that pain induced within the limb performing the motor task, results in increased allocation of attention to that limb, which facilitates motor performance have been discussed [11, 43].

In this study, smoothness of movement improved at a similar rate for all groups across training, although the intervention group was smoother at all time points compared to the control group, whereas the persistent pain group did not differ from the healthy participants regarding smoothness of movement. The latter finding contrasts with research exploring the impact of clinical pain on the smoothness of movement. A common pattern of motor adaptation to pain which has been described in the literature is that pain reduces smoothness of movement [44, 45], alters movement variability and increases co-contraction of muscles resulting in movement stiffness [46]. Recently, individualised motor responses to pain have been reported [47, 48]. Increasingly the impact of pain on smoothness of movement and movement variability are being associated with cognitive aspects of pain such as fear of pain [48] and fear of movement [44] as well as pain intensity.

The results of this study provide a number of observations which provide further information to help understand the relationship between pain, attention and facilitation of performance. Firstly, the increased accuracy and smoothness of movement at all time points was not observed in the pain-intervention group, suggesting that the presence of other factors associated with experiencing persistent pain may modulate the relationship between pain and allocation of attention. Secondly, there was no impact of the ‘movement the pain was associated with’ on the accuracy and smoothness of movement for individual targets, suggesting that increased attention was not specific to the exact movement that caused the pain, providing further insight into the specificity of attention allocation. Previous research has demonstrated that non-noxious electro-cutaneous stimulation to a limb can increase attention to that limb and increase performance [49]. Building on this research, the current study suggests a salient experience that demands attention such as pain is a stronger facilitator of performance than a non-noxious stimulus, in healthy individuals.

### Gaze Behaviour and Attentional Control during Motor Learning

The findings in this study provide support to the processing efficiency theory [7]. The eye gaze behaviour described above may be evidence of higher performance effectiveness, for example, greater use of attentional resources during early training and a decrease in use of attentional resources during later training accompanied by improved performance. The decrease in time to initial orientation of attention to the target in late training aligns with findings from previous studies that show individuals increasingly employ strategies aimed at early seeking of information related to the task, as learning progresses [4, 5]. Safstrom et al. (2014) suggested early attentional allocation to targets was associated with earlier motor action. In the present study, target entry time positively correlated with the time it took to complete sequences in the control group supporting this link between initial orientating and earlier and more efficient movement performance. Despite the early allocation of attention to the next target, the target exit time did not significantly change across training as may be expected. Safstrom et al. (2014) utilised a similar MSL task which required the participants to hold the cursor over the target for a short period of time. The authors reported participants disengagement from the previous target reflected a gaze behaviour that prioritised monitoring the cursor location for the full period of the hold. The findings of the present study support this observation and the idea of a dual role for attention for MSL tasks which require periods of arrest over a stimulus. This provides support for the theory of the development of an attention schema [16] which may be specific to the motor task being employed.

The above pattern of early orientation of gaze to the target and potentially away from the cursor as training progresses has been associated with the move away from a dependency on error-based learning observed in early acquisition [5]. This theory may also underlie the significant reduction in fixation count related to targets observed across training. MSL requires the development of a spatial map of task related stimuli [4, 50]. Therefore, effective attentional engagement and maintenance on a target provides an opportunity to gain information both on the spatial characteristics of targets and the predicted error between said locations and motor action. A significant negative correlation between the above eye gaze indices and both overall motor performance provide support for this theory. This suggests that a more efficient engagement and maintenance of attention of the targets results in better performance. It can therefore be suggested that the number of fixations observed in this study during early training plays a role in facilitating error-based learning processes and building of the spatial map.

Finally, the majority of findings in this study showing significant correlations between eye gaze indices and performance measures, are seen during early training. This may support theories that attentional processes play a larger role in performance improvements during early learning compared to late learning [4]. Whether consciousness is important to the control of these attentional processes is not clear, but the importance of attention during earlier stages may feed discussions around the importance of explicit learning processes during early acquisition of novel MSL.

### The Impact of Pain on Gaze Behaviour and Attentional Control during Motor Learning

Pain is thought to demand attention [8] and is designated as high priority when considering attentional allocation [51]. Despite this, attentional allocation during learning was not significantly different between the two groups receiving experimental pain, and the control group receiving a non-noxious stimulus. This finding contradicts much of the previous research that suggests pain impairs attentional allocation [52–54]. The findings also do not provide specific evidence for pain facilitating attention and therefore motor performance during motor learning, a theory proposed in previous studies exploring pain interference with motor learning [9–13] and discussed above.

The lack of impact of experimental pain on attentional control during motor learning demonstrated in this study, could be due to the motor task and demands of the painful stimuli not exceeding the available attentional resources, as outlined by the ‘biased competition’ model [55]. It is possible that the sequence and subsequent motor actions were not complex enough to require a large enough allocation of attention. Taylor and Thoroughman (2007) reported increasing interference between dual tasks with increasing task complexity, suggesting this was down to the increased attentional requirements of the more complex task. Alternatively, the pain stimuli may not have been assigned higher enough value and the participants in this study may have chosen to prioritise the motor learning tasks over the painful stimulus maintaining their pursuit of the non-pain goal [57] and therefore reducing the attention allocated to the pain. This pattern of attentional allocation has been reported previously and is commonly accompanied by a reduction in the reported levels of pain [57, 58] which was not seen in the present study.

Further evidence of a more nuanced impact of experimental pain on attentional processes was observed in the relationship between psychological constructs, such as pain catastrophizing and fear avoidance, and attention engagement measures in the intervention group. Participants with higher levels of pain catastrophizing had greater numbers of fixations on targets in early training, whereas participants with higher levels of fear avoidance had a lower number of fixations on targets in late training. This provides further support to the theories that state that attentional allocation may depend on factors such as participant motivations and judgements related to value [58]. Interpreting these findings through the lens of the fear avoidance model [59], it could be argued that during initial exposure to the painful stimulus during early acquisition, higher levels of pain catastrophizing results in higher value placed on pain related stimuli, leading to hypervigilance to the stimuli and higher number of visits to the targets. Following a period of learning the participants with higher levels of fear avoidance showed greater adaptation to reduce attentional engagement with the target. This gaze behaviour seen across training in the intervention group in this study has similarities to the Pavlovian learnt behaviours to pain discussed by Meulders (2020). It is important to state that this study provides no evidence of a consistent link between psychological constructs, attention and task performance and therefore the above comments should be seen as exploratory.

## Strengths and Limitations

There are a number of strengths of this present study. Firstly, the study used low frequency sinusoidal electrical stimulation as a way of inducing a valid movement-evoked pain across a sustained period of time which opens up the possibility of exploring the impact of movement relevant phasic pain, commonly seen in musculoskeletal conditions on sustained periods of activity. The use of a non-noxious electro-cutaneous stimulation in the control group provided a method of distinguishing the specific effects resulting from the actual pain sensation from the presence of a non-noxious stimulus that demands attention. This is particularly important as previous research has suggested that allocation of attention is an important mediator of pain interference. Secondly, the use of eye gaze technology to collect data on gaze behaviour, an indirect measure of attention, provided new information on potential mechanisms underlying facilitation of performance in the presence of pain. Finally, the inclusion of a population with a previous experience of persistent pain provided further opportunity to explore factors that influence pain interference. The persistent pain group reported greater levels of pain catastrophizing and fear of pain and inclusion of this population may provide future studies with opportunities to understand the complex interactions between pain, attention and motor learning.

The following limitations of the study need to be considered when interpreting its findings. Although the pain levels were consistent across trials the average pain levels for the intervention group were significantly different from the target level of five out of ten discussed in the methodology. There was also a large range of pain scores for each trial within each group. Pain intensity is associated with greater [52] and longer [60] pain interference and therefore a low level of pain intensity may have influenced the interaction between pain and motor learning. The size of the pain intervention group was small which was exacerbated by the exclusion of some participants due to poor quality of eye gaze data. Due to the small sample results with small effect sizes should be interpreted with caution.

## 5. Conclusion

This study provides initial evidence that low-frequency electro-cutaneous stimulation can be used as a valid method to provide movement-contingent pain across a sustained period of time. The study found that the presence of movement-contingent pain in healthy participants did not impact on improvements in performance across a visuomotor sequence training reflecting the findings of previous research utilising tonic pain paradigms. Similar findings in a group experiencing persistent pain and reporting higher catastrophizing scores provides preliminary evidence that the lack of impact of pain on motor learning is consistent across populations with variable pain experience and attitudes towards pain. Finally, the increased task accuracy at all time points seen in the healthy group experiencing experimental pain, and reported in previous studies, was not associated with different attentional strategies. This finding fails to support theories that increased performance reported throughout motor learning in healthy subjects in the presence of pain is down to improved attention or focus on the task.

## Acknowledgements

The authors would like to thank Helio Cabral and Valter Devecchi for adapting an already established MATLAB programme for the movement-contingent experimental pain paradigm.

## Data availability

The data sets are available from the corresponding author on reasonable request. Please contact: for data enquiries.

## Author contributions

DM, DF, and AK were responsible for the conception of the research question, development of the methodology and drafting of the manuscript. DM completed the data collection and data analysis. All authors have approved the final manuscript and contributed to data interpretation, conclusions and dissemination.

## Supporting information

**S1 Table: History of Pain location and levels for participants in the pain-intervention group.**

**S2 Table: Details of reasons why participants excluded from data analysis.**

**S3 Table: Settings for eye tracking.**

**S1 Fig: Diagram demonstrating how accuracy was calculated.**

